# Optimized chemical labeling method for isolation of 8-oxoG-modified RNA, ChLoRox-Seq, identifies mRNAs enriched in oxidation and transcriptome-wide distribution biases of oxidation events post environmental stress

**DOI:** 10.1101/2024.06.15.599069

**Authors:** Matthew R. Burroughs, Philip J. Sweet, Lydia M. Contreras

**Affiliations:** McKetta Department of Chemical Engineering, UT Austin, Austin, TX 78712; Department of Molecular Biosciences, UT Austin, Austin, TX 78712

**Keywords:** 8-oxoG, RNA Modifications, Epitranscriptome, Air Pollution, Oxidative Stress, Next Generation Sequencing, RNA Oxidation, BEAS-2B Cells, Bioinformatics, Environmental Stress

## Abstract

Bulk increases in nucleobase oxidation, most commonly manifesting as the guanine (G) nucleobase modification 8-oxo-7,8-dihydroguanine (8-oxoG), have been linked to several disease pathologies. Elucidating the effects of RNA oxidation on cellular homeostasis is limited by a lack of effective tools for detecting specific regions modified with 8-oxoG. Building on a previously published method for studying 8-oxoG in DNA, we developed ChLoRox-Seq, which works by covalently functionalizing 8-oxoG sites in RNA with biotin. Importantly, this method enables antibody-free enrichment of 8-oxoG-containing RNA fragments for Next Generation Sequencing-based detection of modified regions transcriptome-wide. We demonstrate the high specificity of ChLoRox-Seq for functionalizing 8-oxoG over unmodified nucleobases in RNA and benchmark this specificity to a commonly used antibody-based approach. Key advantages of ChLoRox-Seq include: (1) heightened resolution of RNA oxidation regions (e.g. exon-level) and (2) lower experimental costs. By applying ChLoRox-Seq to mRNA extracted from human lung epithelial cells (BEAS-2B) after exposure to environmentally relevant stress, we observe that 8-oxoG modifications tend to cluster in regions that are G-rich and within mRNA transcripts possessing longer 5’ UTR and CDS regions. These findings provide new insight into the complex mechanisms that bias the accumulation of RNA oxidation across the transcriptome. Notably, our analysis suggests the possibility that most mRNA oxidation events are probabilistically driven and that mRNAs that possess more favorable intrinsic properties are prone to incur oxidation events at elevated rates. ChLoRox-Seq can be readily applied in future studies to identify regions of elevated RNA oxidation in any cellular model of interest.

## Introduction

Reactive Oxygen Species (ROS) are ubiquitous in nature, manifesting from endogenous sources including mitochondrial processes [1], tumor microenvironments [2], and other disease pathologies [3,4] or exogenous sources such as air pollution [5] and ionizing radiation [6]. The presence of intracellular ROS can prove detrimental to cell fate by directly reacting with fundamental biological molecules (e.g., proteins, lipids, carbohydrates, DNAs, and RNAs), thereby compromising their functionality [7]. Evidence in human cells suggests that oxidative damage accumulates unevenly across the transcriptome, resulting in specific RNA sequences becoming differentially oxidized under oxidative stress [8–11]. These early investigations into the distribution of RNA oxidation have been limited by resolution, evaluating differential accumulation of oxidative damage at the whole transcript level. However, identifying more specific regions (e.g. individual transcript exons) of RNA oxidation accumulation can better serve to prognosticate both (1) the underlying factors that influence where within the transcriptome RNA oxidation occurs and (2) the early impacts of oxidative stress on the cell.

The guanine (G) nucleobase present in DNA and RNA possesses a characteristically low redox potential relative to the other canonical nucleobases [12], lending itself to be a prime target of chemical oxidation by ROS. The most common chemical byproduct formed due to guanine oxidation is 8-oxo-7,8-dihydroguanine (8-oxoG), manifesting in approximately 1 in every 10^5^ G sites at basal levels [13]. Global levels of the free 8-oxoG nucleoside in mediums such as blood, urine, and saliva have been used to assess oxidative stress levels in the human body [14,15]. Additionally, perhaps owing to the mutagenicity of this modification, bulk 8-oxoG levels are upregulated in several chronic human neurological diseases (e.g., Alzheimer’s Disease [8,16,17], Parkinson’s Disease [18] etc.). As such, the 8-oxoG chemical modification has garnered interest for study as a biomarker of oxidative stress. In human DNA, 8-oxodG damage is corrected by the base excision repair protein, 8-oxoguanine DNA glycosylase (OGG1) [19–21]. However, despite RNA being more susceptible to oxidative damage than DNA [22,23], such a mechanism for oxidative damage repair in human RNA remains unknown.

In targeted mechanistic studies, 8-oxoG has been shown to elicit drastic impacts on the fate of RNAs. Seminal work demonstrated that the presence of 8-oxoG within an mRNA transcript increases the propensity of ribosomal stalling at the modification site [24]. This stalling phenomenon has been further shown to result in no-go decay (NGD) processing of modified mRNAs [25]. Additionally, the promiscuous base-pairing capacity of this modification lends itself to tRNA codon misalignment and other mutagenic consequences [26]. While these targeted mechanistic studies reveal possible consequences of 8-oxoG modification to mRNA, methods for detecting locations where these modifications accumulate within specific mRNAs (and the different patterns of modification accumulation that likely arise during oxidative stress) are required to fully understand the impact of RNA oxidation on cellular homeostasis and its contribution to disease progression.

A widely used strategy for performing transcriptome-wide RNA modification mapping has been through an **R**NA-**I**mmuno**P**recipitation and **Seq**uencing (RIP-Seq) approach [27]. In this method, a modification-specific antibody is used to enrich for modification-containing RNA strands (pulldown fraction) from a heterogenous total RNA pool (input fraction). Both pulldown and input RNA fractions are subjected to **N**ext **G**eneration **S**equencing (NGS) analysis, and an increase in the relative abundance of an RNA sequence in the pulldown fraction is used as a proxy for elevated RNA modification abundance on specific RNA transcripts [28–31]. We have previously employed this strategy to investigate the 8-oxoG landscape in mRNA upon environmentally imposed oxidative stress and discovered that some mRNAs are differentially oxidized [9,10]; however, limitations with this method (e.g. high cost, off-target capture, low recovery, etc.) warrant the development of alternative experimental approaches for transcriptome-wide investigation of 8-oxoG accumulation patterns in RNAs.

Antibody-free methods of detection have been established to study 8-oxoG in DNA. Notably, Ding et al. (2017) capitalized on the inherent reactivity of the 8-oxoG modification in their OG-Seq method [32]. In this method, a mild oxidant, K_2_IrBr_6_, is used to specifically generate an unstable intermediate product at 8-oxoG modification sites. This unstable intermediate reacts with a commercially available EZ-link^TM^ Amine-PEG2-Biotin compound, forming a covalent linkage. As a result, an adduct is generated specifically at the 8-oxoG modification site that is functionalized with a biotin tag. The stabilized, biotinylated 8-oxoG adduct is subsequently amenable for affinity-based capture and sequencing. OG-Seq was first applied to investigate the 8-oxoG landscape in DNA isolated from mouse embryonic fibroblasts; this study unveiled that 8-oxoG marks appear to differentially accumulate in gene promoter regions and UTRs [32]. Another study found that DNA promoter regions harboring 8-oxoG modifications in mouse adipose tissue are generally enriched in GC-rich transcription factor binding sites [33]. More recently, a study modified the OG-Seq strategy to investigate the presence of 8-oxoG in miRNAs [34]. However, the 8-oxoG labeling molecule used in this method requires complex organic synthesis and is not commercially available, severely limiting the potential breadth of application. Similar chemistries have also been employed to investigate the spatial occurrence of 8-oxoG with intracellularly localized oxidative stress [35,36]. Yet, such techniques rely on the expression of ectopic engineered proteins and are thus removed from the natural oxidative stress context.

Alternatively, other 8-oxoG detection strategies have utilized the mutagenic 8-oxoG:A Hoogsteen-type base-pairing signature to detect 8-oxoG locations in cDNA reverse transcribed from RNA by monitoring 8-oxoG>T sequencing variations [37–39]. Such a strategy was used to demonstrate the tendency of 8-oxoG modifications to cluster in the seed regions of human miRNAs [38]. While this type of approach provides promise for site-specific resolution of modification sites within RNAs, it is incapable of distinguishing 8-oxoG-induced mutational cDNA signatures from the signatures of other G-derived RNA modifications that include 5-guanidinohydantoin (Gh) and spiroiminodihydantoin (Sp). As a result, interpretation of true positive 8-oxoG sites is difficult.

In this study, we present a new, antibody-free strategy for profiling the location of 8-oxoG modifications in mRNA at the transcriptome-wide level—**Ch**emical **L**abeling **o**f **R**NA targeting 8-**ox**oG modifications and **Seq**uencing (ChLoRox-Seq). For this method, we optimized the chemical labeling reaction conditions outlined in OG-Seq for DNA [32] to covalently attach a biotin moiety to 8-oxoG modification sites in RNA with high specificity, thereby enabling biotin-centric detection and streptavidin-mediated enrichment. We further demonstrated the high specificity of this optimized labeling reaction for 8-oxoG by benchmarking against an anti-8-oxoG antibody used in conventional RIP-Seq analysis. We subsequently applied ChLoRox-Seq to investigate the transcriptome-wide distribution of 8-oxoG sites at the individual transcript exon level in a model of environmentally induced oxidative stress. We exposed human lung cells (BEAS-2B) to liquid suspensions of two well-documented sources of oxidative stress: **P**articulate **M**atter (PM) [40,41] and hydrogen peroxide (H2O2) [42]. Notably, we discovered a population of mRNAs demonstrating extreme susceptibility to RNA oxidation—identified as having oxidized exon regions across all tested exposure conditions. From this population of readily oxidized mRNAs, we identified intrinsic characteristics that appear to dictate mRNA susceptibility to oxidative damage. Our findings demonstrate a robust, cost-effective strategy for detecting the location of 8-oxoG modifications in mRNA with single exon resolution that can be easily adapted to investigate the 8-oxoG modification landscape in any species of interest.

## Results

### Optimized biotin labeling reactions accomplish specific labeling of 8-oxoG in RNA without degrading RNA

Prior work demonstrated that, under mild redox conditions using a one-electron oxidant (K_2_IrBr_6_), 8-oxoG modifications in DNA can be functionalized with an amine-terminated biotin linker molecule [32] (Fig. 1, A). However, when directly applying these reaction conditions to total RNA isolated from K-12 wild-type *Escherichia coli* (*E. coli*), we found that these conditions resulted in significant sample degradation (Fig. 1, B). We hypothesized that the high-temperature specifications (75°C, 30 minutes), combined with elevated concentrations of chemical oxidant and labeling molecules, were likely contributing to this observed degradation. By reducing temperature and reactant concentrations, we preserved RNA integrity (Fig. 1, B). Next, we sought to optimize the specificity of this biotin labeling reaction to target 8-oxoG modifications over unmodified RNA. To investigate reaction specificity, we applied a dot-blot experimental approach (Fig. 1, C). In these experiments, we captured chemiluminescent signal-based comparison of biotin incorporation into short (24-25-nt) synthetic RNA sequences that were commercially synthesized to either contain the 8-oxoG modification (in the 19^th^ position) or were unmodified. We computed a reaction specificity ratio by dividing the chemiluminescent signal output from 8-oxoG-containing RNAs by the signal from unmodified RNAs for labeling conditions across a gradient of reaction temperatures. We found that room temperature (RT, ∼20°C) conferred a marked increase in specificity (approximately 78-fold) compared to elevated reaction temperatures (i.e. approximately 14-fold at 49°C) (Fig. 1, D-E). Ultimately, we arrived at labeling reaction conditions that generated optimal specificity towards 8-oxoG-modifed RNA over unmodified RNA (Table 1). These conditions were used in all subsequent biotin labeling reactions referenced throughout this work.

**Figure 1.**
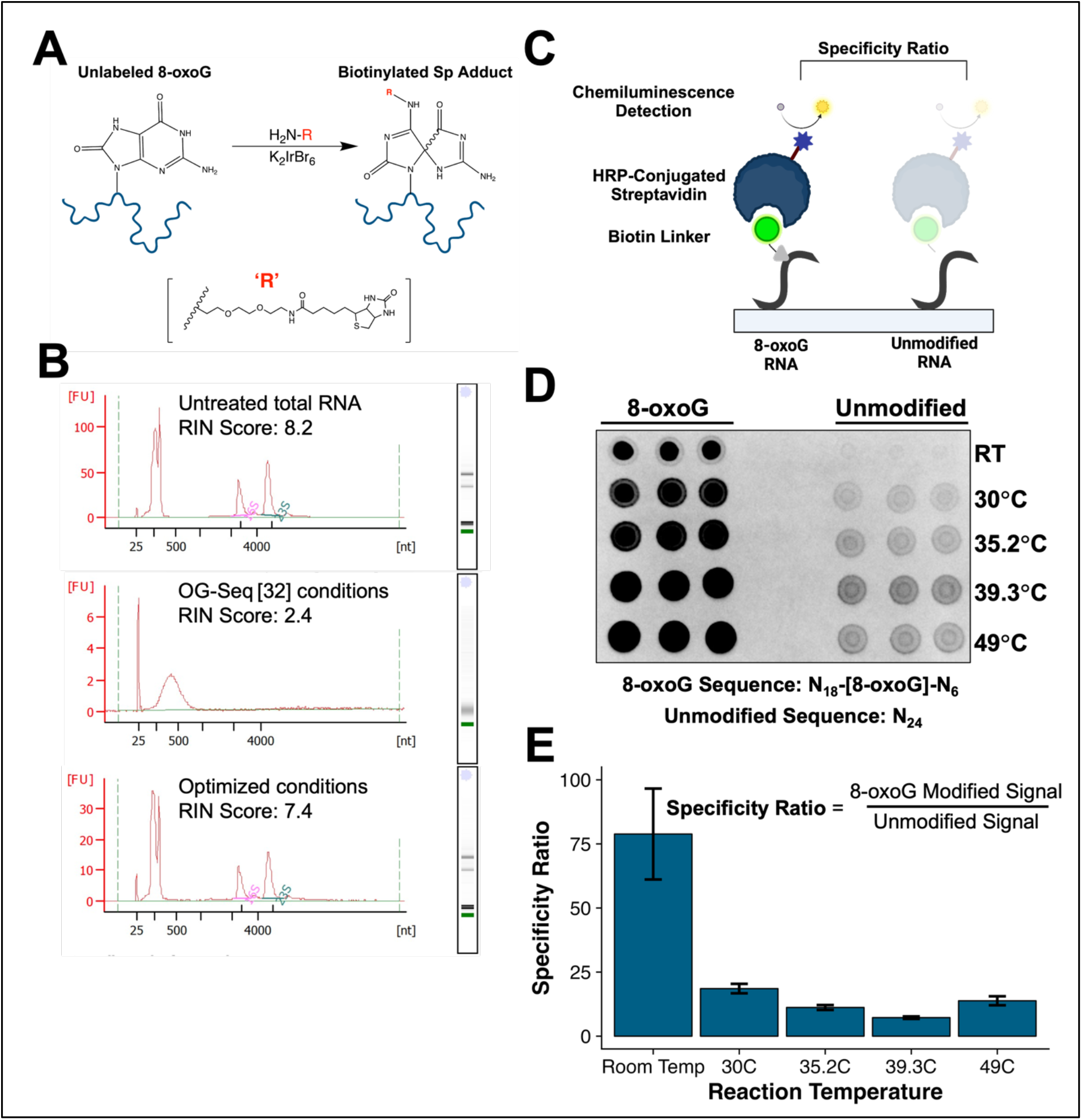
Optimized biotin labeling reaction conditions confer specific labeling of 8-oxoG without degrading RNA. (A) Schematic of labeling procedure used to functionalize 8-oxoG modifications with biotin in RNA. (B) Agilent Bioanalyzer 2100 electropherograms of *E. coli* total RNA untreated, labeled using OG-Seq reaction conditions, and labeled using optimized reaction conditions. (C) Schematic of dot blot assay used to evaluate biotin labeling specificity for 8-oxoG-modified RNA over unmodified RNA. (D) Dot blot image of biotin labeling reaction products for 8-oxoG modified RNA versus unmodified RNA over a temperature gradient. (E) Bar graph of specificity factors (8-oxoG signal / Unmodified signal) for the biotin labeling reaction at each temperature quantified using Bio-Rad ImageLab™ software.

**Table 1.** Optimized biotin labeling reaction conditions used in ChLoRox-Seq compared to conditions used in OG-Seq [32].

| Reaction Component | ChLoRox-Seq Quantity | OG-Seq [32] Quantity |
| --- | --- | --- |
| Time | 30 minutes | 30 minutes |
| Temperature | Room Temperature (20°C) | 75°C |
| RNA/DNA Input | Variable | Variable |
| [K2IrBr6] | 0.5 mM | 5 mM |
| [EZ-link™ Amine-PEG2-Biotin] | 2.5 mM | 20 mM |
| [NaPi Buffer] | 100 mM | 100 mM |
| Buffer pH | 8.0 | 8.0 |
| RNase Inhibitor (Murine) | 0.4 U/μL | -- |

### The biotin labeling approach offers comparable specificity towards 8-oxoG as the anti-8-oxoG antibody

We next sought to benchmark the specificity of biotin labeling 8-oxoG against an anti-8-oxoG antibody previously used for RIP-Seq detection methodologies [9,10]. Conveniently, antibodies are amenable to specificity analysis using an analogous dot blot scheme. Therefore, we used the same dot blot approach to compare the specificity of the 8-oxoG antibody and the biotin labeling reaction (Fig. 2, A). We found that the specificity of the biotin labeling reaction was comparable to that of the 8-oxoG antibody (Fig. 2, B), with the biotin labeling reaction generating a mean specificity ratio of approximately 94-fold and the anti-8-oxoG antibody generating a mean specificity ratio of approximately 67-fold. We further tested cross-reactivity for the biotin labeling reaction towards labeling another oxidation-related purine RNA modification, 8-oxo-7,8-dihydroadenine (8-oxoA) [43]. Despite the structural similarity between 8-oxoA and 8-oxoG, as well as their similar redox potentials [12], we noted a relatively low, approximately 2-fold labeling preference for 8-oxoA over unmodified RNA (Supp. Fig. S1).

**Figure 2.**
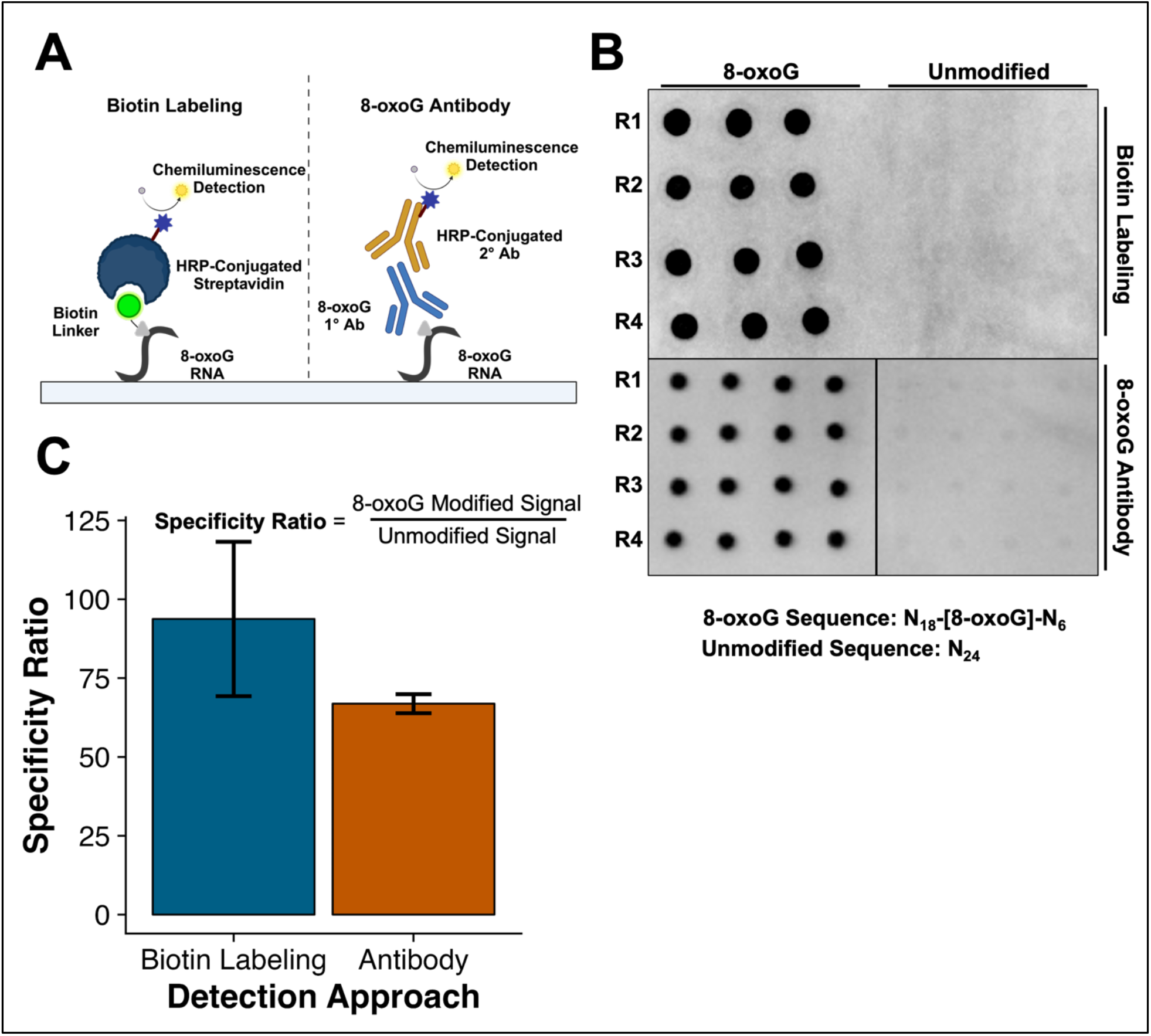
The biotin labeling approach offers comparable specificity towards 8-oxoG as the anti-8-oxoG antibody. (A) Schematic of dot blot method for comparing 8-oxoG detection specificity between biotin labeling approach and antibody-based approach. (B) Dot blot image of room temperature biotin labeling detection versus antibody-based detection for 8-oxoG modified RNA versus unmodified RNA. R1-R4 indicates independent technical replicates. (C) Bar graph of specificity factors for each detection method quantified using Bio-Rad ImageLab™ software.

### The biotin labeling approach captures increasing bulk 8-oxoG levels in total RNA exposed to increasing oxidative stress in vitro and in vivo

Upon establishing that the biotin labeling reaction was specific towards the 8-oxoG modification over unmodified RNA, we tested if we could capture increases in the relative abundance of 8-oxoG in a diverse pool of total RNA exposed to elevated ROS conditions. First, we *in vitro* exposed total RNA isolated from K-12 wild-type *E. coli* to increasing levels of ROS using Fenton chemistry with hydrogen peroxide (H2O2) and Fe^2+^, as outlined in previous work [31]. We used the same dot blot method to evaluate increases in bulk 8-oxoG modifications in the samples and compared between biotin labeling and antibody-based 8-oxoG detection methods. We found that both strategies were amenable for detecting statistically significant increases in 8-oxoG abundance at elevated ROS conditions (Fig. 3, A-B).

**Figure 3.**
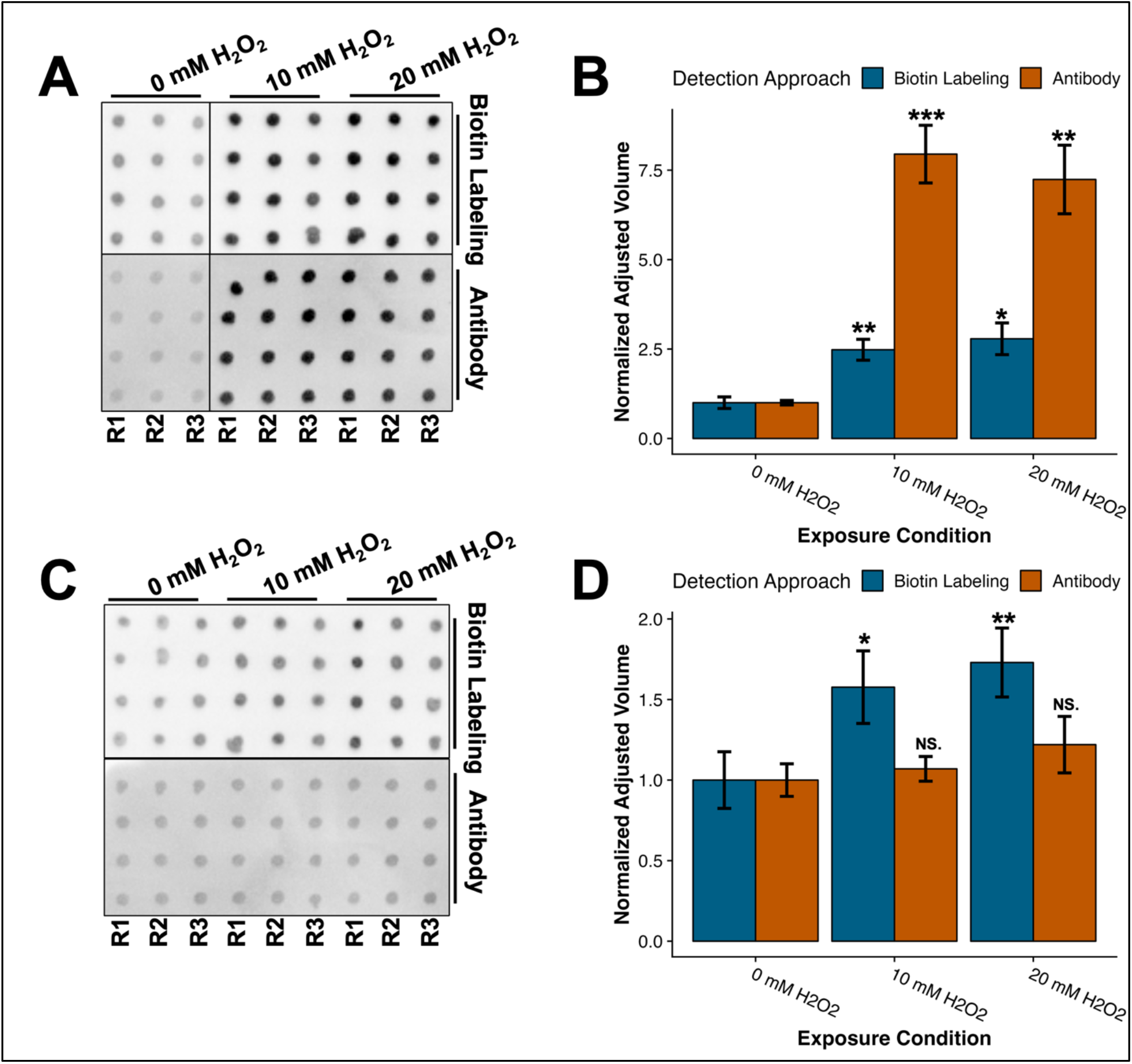
The biotin labeling approach captures increasing 8-oxoG levels in total RNA exposed to increasing oxidative stress *in vitro* and *in vivo.* (A) Antibody and biotin labeling dot blot images of *E. coli* total RNA oxidized *in vitro* using Fenton chemistry over a range of increasing H_2_O_2_ concentrations for 1 hour, 37°C. R1-R3 represent independent Fenton reaction replicates. (B) Bar graph of quantified dot blot intensities from (A) using Bio-Rad ImageLab™ software. Normalized Adjusted Volume represents the pixel density of each sample relative to the control, 0 mM H2O2 sample. (C) Antibody and biotin labeling dot blot images of *E. coli* total RNA oxidized *in vivo* over a range of increasing H_2_O_2_ concentrations for 20 minutes, 37°C. R1-R3 represent independent biological exposure replicates. (D) Bar graph of quantified dot blot intensities from (C) using Bio-Rad ImageLab™ software. Normalized Adjusted Volume represents the pixel density of each sample relative to the control, 0 mM H2O2 sample. Data are presented as mean values ± s.d.; NS., *P* ≥ 0.05; (*), *P* < 0.05; (**), P < 0.01; (***), P < 0.001 (Student’s unpaired T-test) relative to the 0 mM H2O2 condition.

We subsequently applied a similar strategy to examine bulk increases in 8-oxoG abundance *in vivo*. Relative to *in vitro* conditions, we expected that the natural protective mechanisms of the cells would contribute to more realistic (i.e. likely lower) levels of RNA oxidation accumulation [43]. Herein, we cultured K-12 wildtype *E. coli* to mid-log phase (OD_600_ = 0.5-0.6) in LB media and then added increasing concentrations of H2O2 (0-20 mM) to induce oxidative stress for 20 minutes at 37°C; these conditions were selected based on previous work in our lab (unpublished data) which found that 20 mM H2O2 exposure at 37°C for 20 minutes resulted in 40-60% survival for K-12. After harvesting cell pellets, we extracted the total RNA as input for dot blot analysis. Encouragingly, we found that the biotin labeling reaction detected statistically significant increases in bulk 8-oxoG abundance whereas the anti-8-oxoG antibody failed to detect any statistically significant increase in 8-oxoG abundance (Fig. 3, C-D). This result validated our biotin labeling reaction as a sufficiently sensitive approach for detecting biologically relevant intracellular accumulation of 8-oxoG modifications.

### Mild exposure conditions generate elevated intracellular ROS levels with insignificant bulk 8-oxoG increases in human BEAS-2B lung cell line

Owing to the mounting evidence of RNA oxidation in human disease [44–46], we were interested in applying this method for the detection of the transcriptome-wide landscape of 8-oxoG modifications in a human model of oxidative stress. Building on previous work in our lab [47], we mimicked lung cell exposure to air pollution by incubating a noncancerous, immortalized human lung cell line (BEAS-2B) in growth media supplemented with two different types of common air pollutants (urban particulate matter (PM) and formaldehyde (FA)). We were interested in selecting experimental exposure conditions under which the intracellular levels of ROS were elevated, but prior to the activation of a transcriptional cellular response that would potentially mitigate the induced accumulation of RNA oxidation. To develop appropriate experimental conditions, we first tracked the accumulation of intracellular ROS in cells exposed to 100 µM H2O2 (H2O2), 500 µM formaldehyde (FA), and 500 µg/mL urban particulate matter (PM) using a fluorescence dye assay, H_2_DCFDA. An unexposed, media-only control (CTRL) was used as a baseline reference of intracellular ROS (Fig. 4, A). We found that, after 20 minutes of exposure at 37°C, 5% CO2, intracellular ROS was elevated for both H2O2 and PM samples, but not for FA samples (Fig. 4, B). Despite elevated ROS at this timepoint, we did not observe significant bulk increases in the level of 8-oxoG modifications in cells exposed to 100 µM H2O2 when evaluated using a biotin dot blot assay (Fig. 4, C-D). It is important to note that bulk increases in 8-oxoG modification levels were not desired, given that we sought to investigate subtle changes to the 8-oxoG landscape under stress. Therefore, exposure conditions of 500 µg/mL urban particulate matter (PM), 100 µM H2O2 (H2O2), and media-only control (CTRL) were selected for use in subsequent ChLoRox-Seq assays to detect regions of 8-oxoG accumulation across the transcriptome.

**Figure 4.**
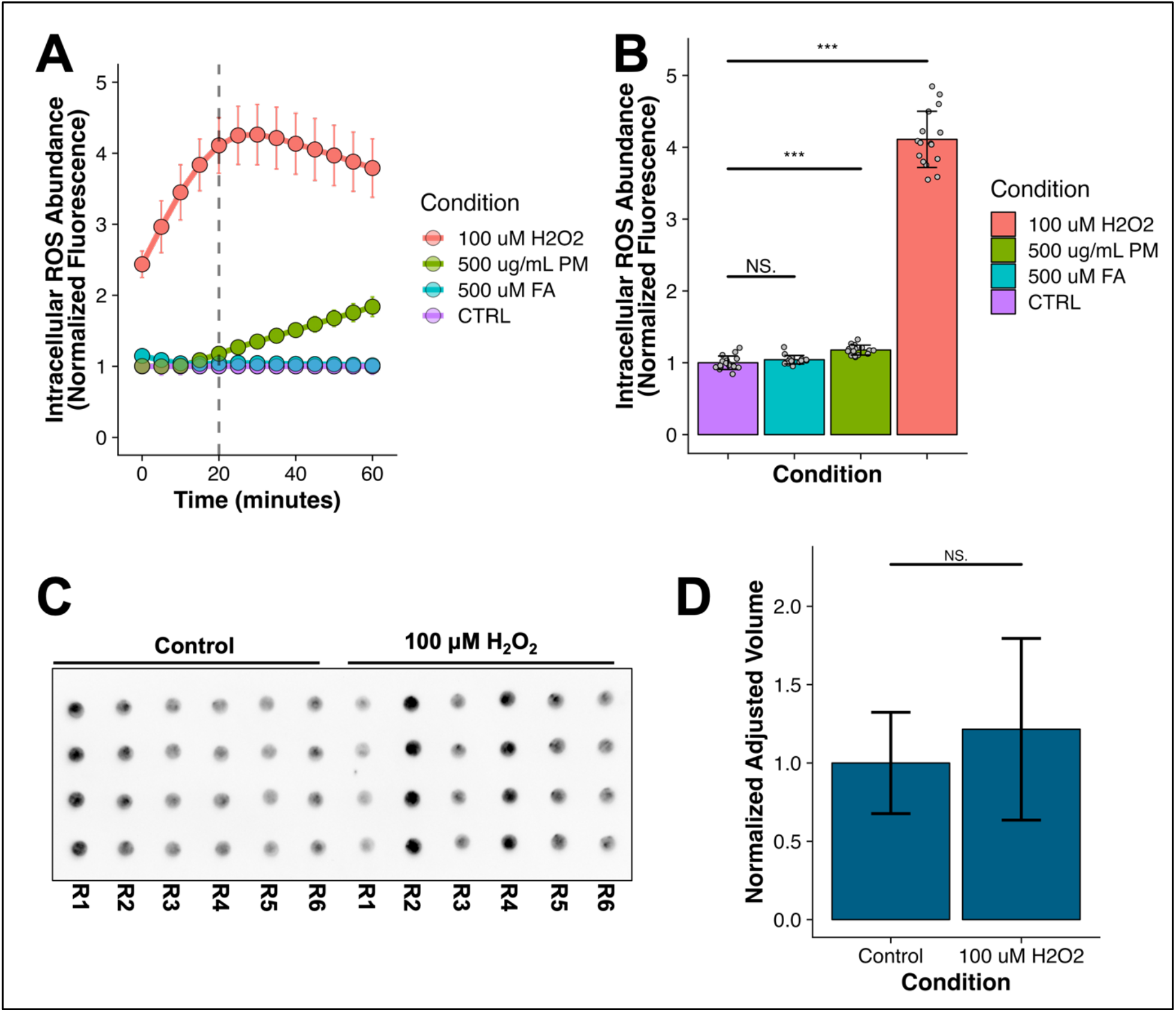
Mild exposure conditions generate elevated intracellular ROS levels with insignificant bulk 8-oxoG increases in human BEAS-2B cell line. (A) H_2_DCFDA assay measuring intracellular ROS accumulation in BEAS-2B cells exposed to CTRL, 500 μM FA, 500 μg/mL PM, and 100 μM H2O2 conditions over a 60-minute time course. Vertical dashed line represents timepoint at which samples were collected for biotin pulldown and sequencing experiment (N = 16). (B) Bar graph of control-normalized fluorescence for each exposure condition after 20 minutes. Gray dots indicate individual biological replicate measurements. (C) Biotin labeling dot blot image of BEAS-2B total RNA following *in vivo* exposure to control media or 100 μM H_2_O_2_ supplemented media for 20 minutes, 37°C. R1-R6 represents independent biological exposure replicates. (D) Bar graph of quantified dot blot intensities from (C) using Bio-Rad ImageLab™ software. Normalized Adjusted Volume represents the pixel density of each sample relative to the control, 0 μM H2O2 sample. Data are presented as mean values ± s.d.; NS., *P* ≥ 0.05; \**P* < 0.05; \*\**P* < 0.01; \*\*\**P* < 0.001 (Student’s unpaired T-test) relative to the control unexposed condition.

### ChLoRox-Seq strategy identifies a select pool of mRNAs that are differentially prone to oxidation

After demonstrating successful selective functionalization of 8-oxoG with biotin in RNA and determining environmental exposure conditions that generate elevated intracellular ROS, we next sought to develop an experimental pipeline for investigating the 8-oxoG modification landscape in a transcriptome-wide fashion. We accomplished this task by developing **Ch**emical **L**abeling **o**f **R**NA targeting 8-**ox**oG modifications and **Seq**uencing or ChLoRox-Seq (Fig. 5, A). See *ChLoRox-Seq Pipeline* in Methods for experimental details. To establish a robust downstream analysis pipeline for this unique isolation method, we subsequently employed three separate bioinformatics approaches to process the Next Generation Sequencing data from this pulldown experiment: DESeq2 [48] with binning reads to individual transcript exon regions, exomePeak2 [49] with GC bias correction, and exomePeak2 without GC bias correction. These three different approaches were selected given that they each pose advantages and disadvantages for analysis of this data type. DESeq2 is a tool widely used for assessing differential expression of read count features (e.g., genes, exons) from RNA-Seq data outputs. exomePeak2 is a peak calling algorithm originally developed for identifying m^6^A modification peaks in RIP-Seq data. For each bioinformatic approach, regions with a log2FoldChange (Pulldown/Input) > 1 and FDR/p_adj_ < 0.05 were classified as oxidized.

**Figure 5.**
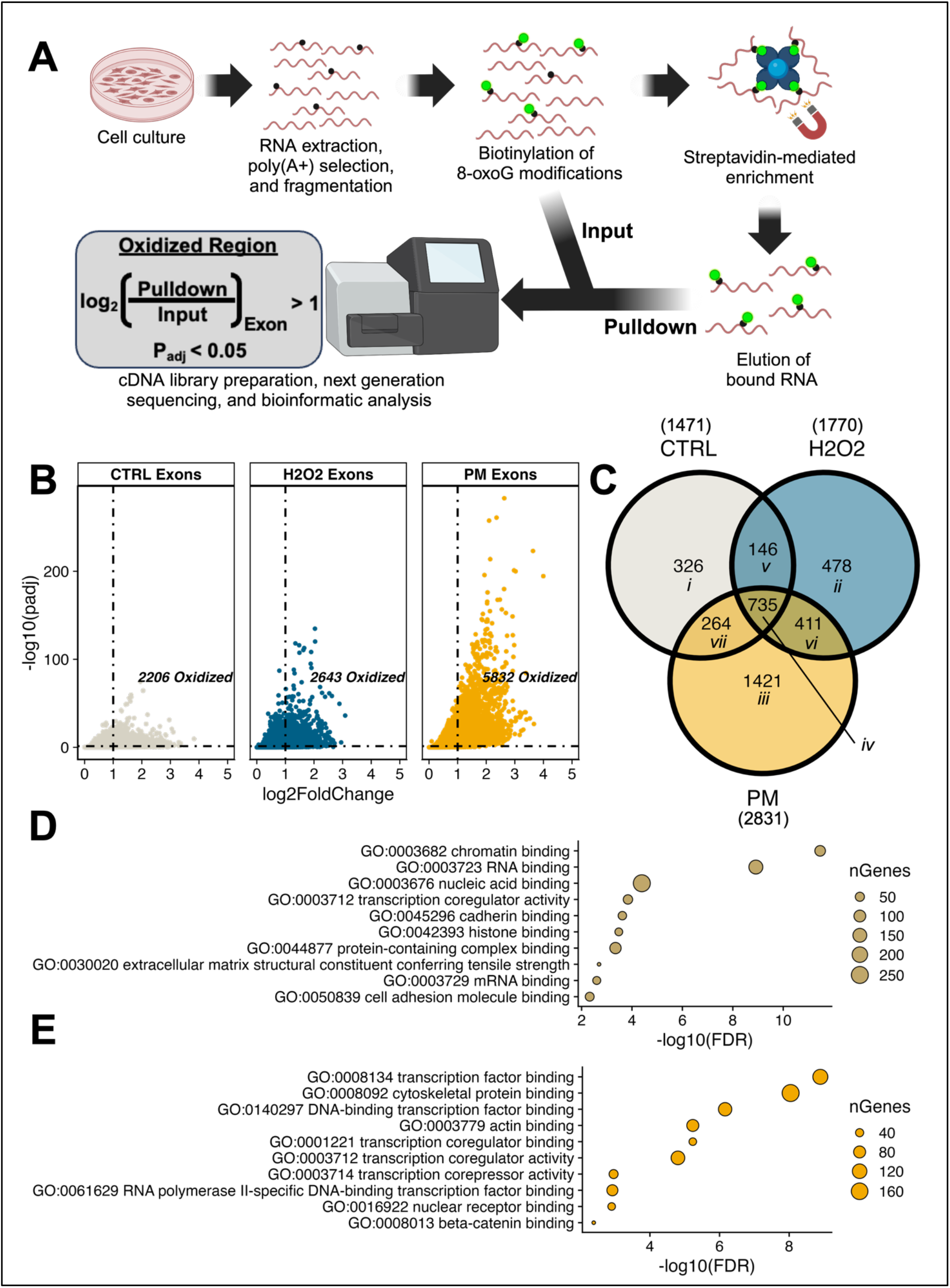
Biotin labeling and pulldown of 8-oxoG-enriched RNA identifies select pools of mRNAs that are prone to oxidation under control and environmental stress conditions. (A) Schematic of ChLoRox-Seq pipeline for identifying regions of RNA oxidation. (B) Volcano plots for each experimental condition depicting exon regions of elevated 8-oxoG abundance. (C) Venn diagram showing overlapping genes containing oxidized regions across each experimental condition. (D) Bubble blot showing top 10 enriched (FDR) GO Molecular Function terms for oxidized genes found across all experimental conditions. (E) Bubble plot showing top 10 enriched (FDR) GO Molecular Function terms for oxidized genes found under PM exposure conditions and not CTRL conditions.

We first investigated oxidation region-containing genes (oxidized genes) that were identified by all three bioinformatics approaches for each of the exposure conditions tested (e.g. CTRL, PM, and H2O2) (Supp. Fig. S3, A). Interestingly, a low degree of overlap was observed between oxidized genes identified using each bioinformatic method (summarized in Table 2). This low degree of overlap was most notable for CTRL (media-only exposed) samples, with a drastic change in the number of oxidized genes observed between DESeq2 with exon binning and exomePeak2 approaches (Table 2, CTRL N=3). To test the hypothesis that this stark difference might result from biological replicate variability, we performed a principal component analysis (PCA) of exon feature read counts across all samples (Supp. Fig. S2, A). Upon doing so, we found that there was indeed a large degree of variation present in one of the CTRL pulldown samples (Supp. Fig. S2, B). Upon removal of the biological pulldown replicate with the highest PCA variability (Supp. Fig. S2, B-D) from analysis, we observed a greater fold increase in the number of identified oxidized region-containing genes for N = 2 compared to N = 3 for both exomePeak2 approaches than the DESeq2 with exon binning approach (Table 2, N = 2 versus N = 3). Unsurprisingly, this observation was most apparent when evaluating CTRL samples, with an approximate 6-12-fold increase in the number of oxidized genes obtained using the exomePeak2 approach compared to an approximate 1.5-fold increase for the DESeq2 with exon binning approach (Table 2, CTRL N = 2 versus N = 3). Based on the extreme dependence of oxidation region detection on biological replicate variability observed for the exomePeak2 approaches, we ultimately decided to proceed with analysis using the DESeq2 with exon binning strategy.

**Table 2.** Overlapping genes containing RNA oxidation regions identified using three different bioinformatics processing pipelines: DE-Seq2 with exon binning, exomePeak2 (+) GC bias correction, exomePeak2 (-) GC bias correction.

| **Exposure Type** | **Pulldown Replicates (N)** | **DESeq2 with exon binning**<br>**($p_{adj} < 0.05$, $L2FC > 1$)** | **exomePeak2 (+) GC bias correction**<br>**(FDR < 0.05, $L2FC > 1$)** | **exomePeak2 (-) GC bias correction**<br>**(FDR < 0.05, $L2FC > 1$)** | **# Conserved** |
| --- | --- | --- | --- | --- | --- |
| **CTRL** | 2 | 2162 | 972 | 1004 | **440** |
|  | 3 | 1471 | 80 | 161 | **42** |
| **# Conserved** | -- | **1197** | **69** | **151** | -- |
| **H2O2** | 2 | 2236 | 1348 | 1336 | **588** |
|  | 3 | 1770 | 672 | 718 | **313** |
| **# Conserved** | -- | **1393** | **530** | **561** | -- |
| **PM** | 2 | 3113 | 5174 | 4936 | **2107** |
|  | 3 | 2831 | 2493 | 2591 | **1313** |
| **# Conserved** | -- | **2551** | **2483** | **2587** | -- |

Using the DESeq2 with exon binning approach, we identified 2206 oxidized exons in CTRL samples, 2643 oxidized exons in H2O2 samples, and 5832 oxidized exons in PM samples (Fig. 5, B; Supp. Fig. S3, B). These oxidized exons belonged to 1471 genes in CTRL samples, 1770 genes in H2O2 samples, and 2831 genes in the PM samples (Fig. 5, C). Many of these genes containing oxidized regions overlapped across all experimental conditions (735), suggesting conservation of RNA oxidation for a select pool of mRNAs. Using the ShinyGO version 0.80 [50] Molecular Function **G**ene **O**ntology (GO) analysis tool, we found that chromatin binding (GO:0003682) and RNA binding (GO:0003723) were significantly enriched functions (FDR < 0.01) in this conserved gene pool (Fig. 5, D). Similarly, we investigated the population of genes that were oxidized in PM samples and not in CTRL samples (Fig. 5, C regions *iii* and *vi*) and found that molecular functions related to transcription factor binding (GO:008134) and cellular structural integrity (GO:0008092, GO:0003779) were significantly enriched (Fig. 5, E). When looking at oxidized genes found in H2O2 and not CTRL (Fig. 5, C regions *ii* and *vi*), we did not observe any molecular function GO terms that passed our significance threshold (FDR < 0.01). A list of the genes present in each subsection of the Venn diagram in Fig. 5, C is included in supplemental table S19.

### mRNAs enriched in the oxidized pool have unique intrinsic properties relative to the general cellular mRNA pool

We next hypothesized that overarching intrinsic mRNA sequence factors contribute to the oxidation susceptibility of specific pools of mRNAs. To test this hypothesis, we utilized general sequence properties obtained from ShinyGO version 0.80 [50] for the pool of oxidized genes conserved across every tested experimental condition (Fig. 5, C region *iv*). We compared these results to the sequence properties for all genes detected in the general (Input) pool of the RNA-Seq data. We found that, on average, mRNAs that were enriched in the Pulldown (Oxidized) pool are longer than the general (Input) pool, with an observed median increase in transcript length from 1692 nucleotides (IQR: 1195 – 2514 nucleotides) to 1819.5 nucleotides (IQR: 1296.5 – 2648.25 nucleotides) (Fig. 6, A). This trend held when evaluating 5’ UTR (Fig. 6, B) and CDS (Fig. 6, C) sequence lengths, with median increases observed from 111 nucleotides (IQR: 74 – 171 nucleotides) to 129 nucleotides (IQR: 86.25 – 198.5 nucleotides) and 897 nucleotides (IQR: 585 – 1424 nucleotides) to 1084.5 nucleotides (IQR: 683.75 – 1807.5 nucleotides), respectively. Interestingly, the 3’ UTR length of mRNAs enriched in the oxidized pool was not significantly different than the general (Input) mRNA pool (Fig. 6, D.). Lastly, the number of exons for mRNAs enriched in the oxidized pool was elevated over the general (Input) pool (Fig. 6, E), with an observed median increase from 6 to 7 exons per transcript; this result was consistent with the fact that mRNAs enriched in the oxidized pool tend to be longer in total sequence length.

**Figure 6.**
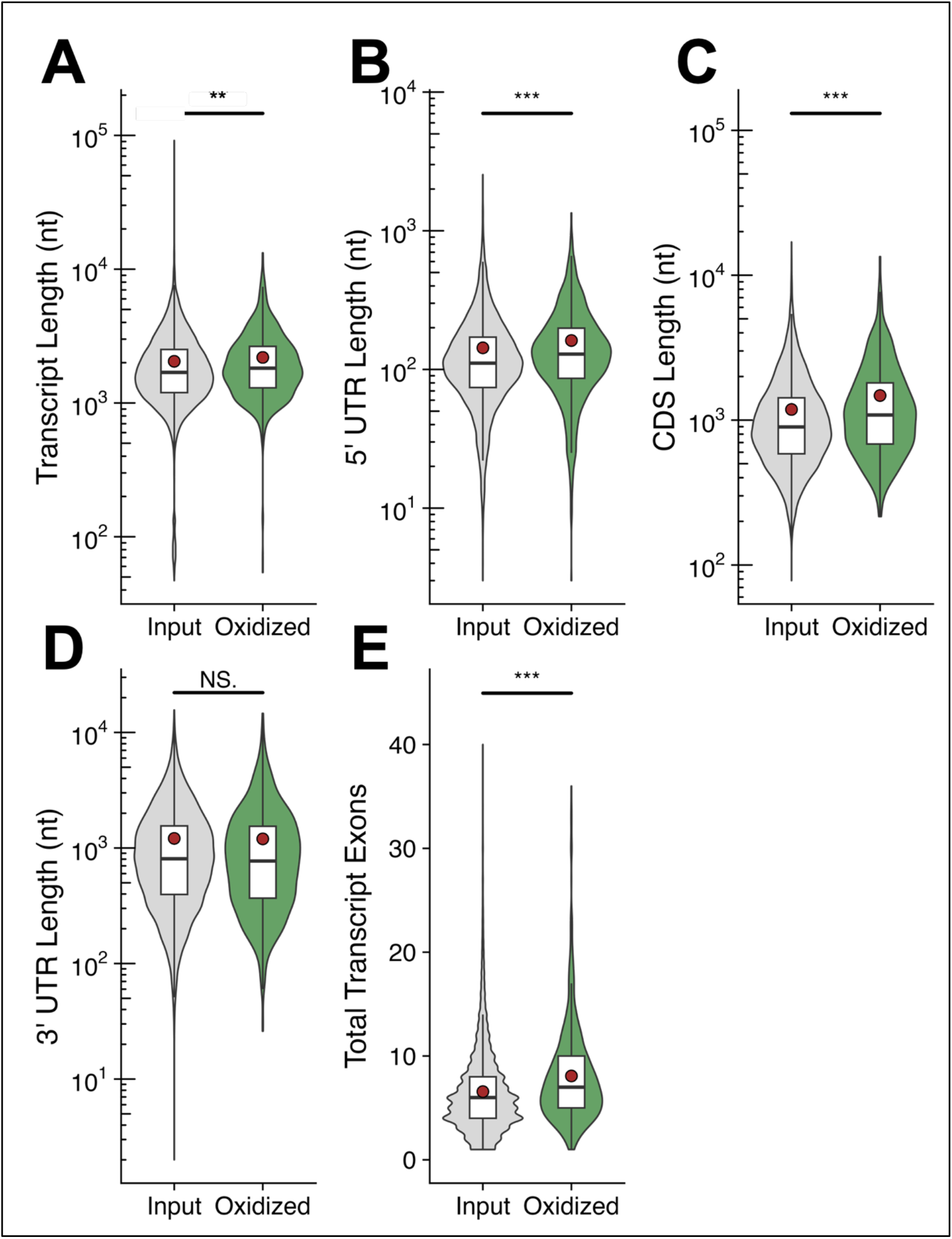
Conserved oxidized genes exhibit unique compositional properties relative to background genes. Boxplots overlaid onto violin plots showing the distribution of (A) transcript lengths, (B) 5’ UTR lengths, (C) CDS lengths, (D) 3’ UTR lengths, and (E) number of exons for Pulldown-enriched mRNAs (Oxidized) versus all mRNAs detected in the Input fraction (Input). Brown circles denote the mean value of the distribution. NS., *P* ≥ 0.05; \**P* < 0.05; \*\**P* < 0.01; \*\*\**P* < 0.001 (Student’s unpaired T-test) relative to the Input fraction. Outlier data points have been excluded from each plot.

### Exons enriched in the oxidized pool are longer and drastically elevated in G nucleotide percentage relative to the input cellular exon pool

Given that the DESeq2 with exon binning bioinformatic approach of ChLoRox-Seq provides exon-level resolution for regions of RNA oxidation—an improvement over previous RIP-Seq experiments, which enriched for whole transcripts [9,10]—we decided to investigate the individual nucleotide composition of exons enriched in the oxidized pool versus the total composition of all exons present in the general (Input) pool of the RNA-Seq data. We found that the percentage of G and C nucleotides was elevated in exons enriched in the oxidized pool relative to exons in the general (Input) pool, with median G percentage increasing drastically from 25.3 % (IQR: 21.3 – 30.1 %) to 39.9 % (IQR: 33.3 – 44.8 %) and median C percentage increasing from 23.5 % (IQR: 18.8 – 29.2 %) to 26.5 % (IQR: 20.9 – 31.1 %) (Fig, 7, A). Correspondingly, the percentage of A and U nucleotides was diminished for exons enriched in the oxidized pool over the general (Input) exon pool, with median A percentage decreasing from 26.6 % (IQR: 21.0 – 31.8 %) to 19.1 % (IQR: 14.1 – 25.7 %) and median U percentage decreasing from 23.0 % (IQR: 18.4 – 27.6 %) to 13.4 % (IQR: 10.0 – 17.8 %) (Fig. 7, A). We additionally observed that exons enriched in the oxidized pool were longer than exons in the general (Input) pool, with a median increase in length from 136 nucleotides (IQR: 95 – 207 nucleotides) to 178 nucleotides (IQR: 105 – 345 nucleotides) (Fig. 7, B).

**Figure 7.**
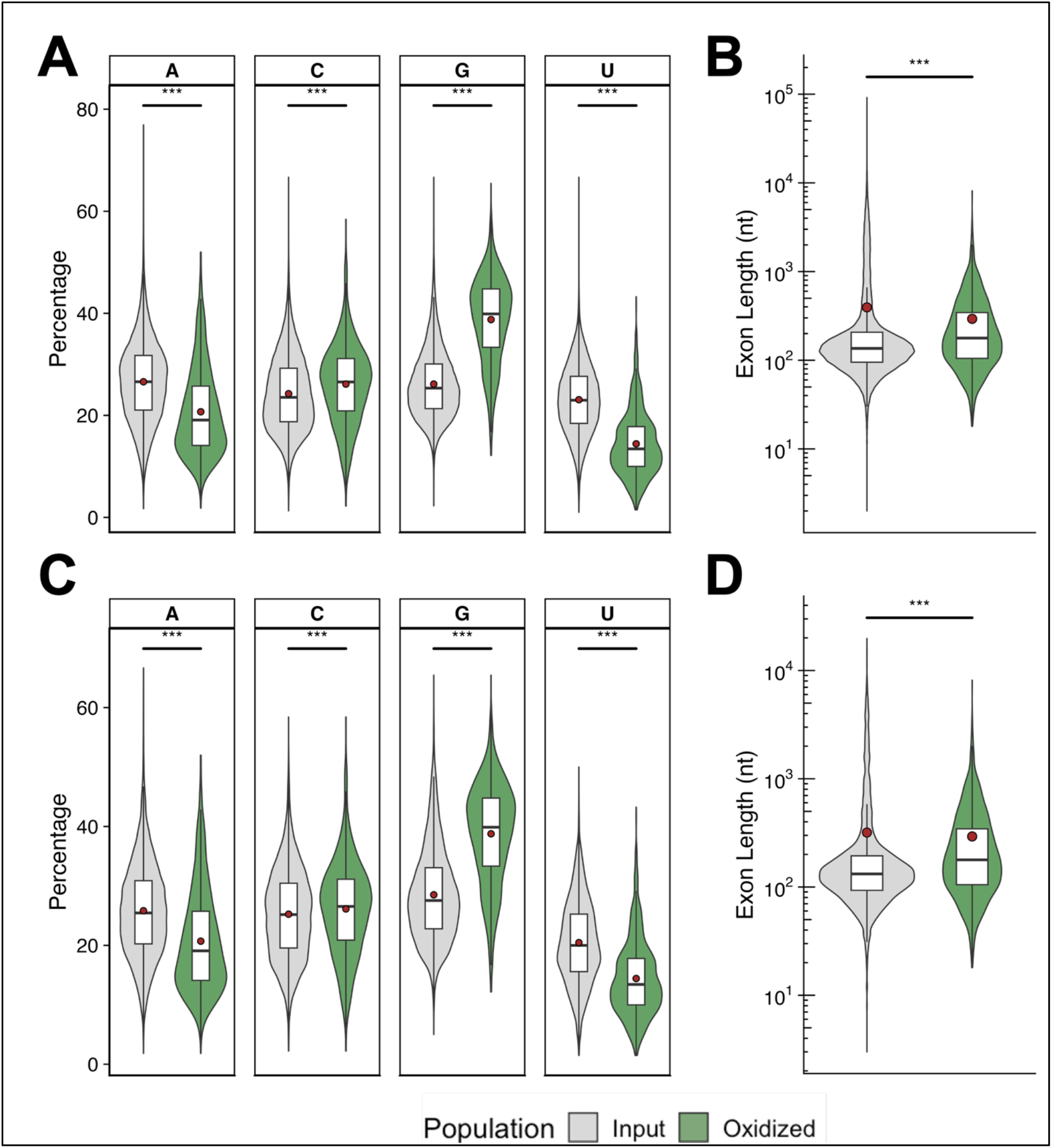
Differentially oxidized exons are longer and enriched in G nucleobases when compared to non-oxidized exons. (A) Distribution of canonical nucleobase contents for exons present in the Input versus Pulldown-enriched (Oxidized) fractions. (B) Distribution of exon lengths for exons present in the Input versus Pulldown-enriched (Oxidized) fractions. (C) Distribution of canonical nucleobase contents for exons present in the Input versus Pulldown-enriched (Oxidized) fractions when restricting the Input to only include exons within genes harboring an oxidized exon. (D) Distribution of exon lengths for exons present in the Input versus Pulldown-enriched (Oxidized) fractions when restricting the Input to only include exons within genes harboring an oxidized exon. Brown circles denote the mean value of the distribution. NS., *P* ≥ 0.05; \**P* < 0.05; \*\**P* < 0.01; \*\*\**P* < 0.001 (Student’s unpaired T-test) relative to the Input fraction. Outlier data points have been excluded from each plot.

We further investigated whether exons enriched in the oxidized pool possess unique properties relative to exons in the general (Input) pool belonging to the same transcript. To guide this analysis, we restricted the number of exons detected within the general (Input) pool to only include exons that are part of genes that contained at least one exon enriched in the oxidized pool, omitting all exons corresponding to genes that were not enriched in the oxidized pool. Again, we observed the same trends as with the total general (Input) exon pool comparison. Exons enriched in the oxidized pool were elevated in G and C residues and depleted in A and U residues relative to other exons on the same transcript identified in the general (Input) pool (Fig. 7, C). Additionally, these exons enriched in the oxidized pool are longer than other exons on the same transcript present in the general (Input) pool (Fig. 7, D).

## Discussion

### ChLoRox-Seq enables the detection and quantification of oxidized RNAs

RNA oxidation is of keen interest in the medical community, being intimately linked to several oxidative stress-related health disorders [44–46]. 8-oxoG is the most abundant RNA modification associated with cellular exposure to ROS. While targeted mechanistic studies offer a glimpse into the myriad impacts of 8-oxoG on RNA, a lack of available whole-transcriptome methods for identifying the location of this modification has prevented a holistic understanding of the underlying factors that bias the distribution of this modification. In this work, we have developed a new, cost-effective, species-agnostic method for determining the location of 8-oxoG modifications in RNA. In this method, we functionalize 8-oxoG residues with biotin using an optimized redox chemistry-based strategy motivated by a previous study investigating 8-oxoG in DNA (Fig. 1, A). We showcase how biotinylated 8-oxoG residues are subsequently amenable for streptavidin-based affinity capture and NGS analysis to determine the transcriptome-wide distribution of 8-oxoG modifications at the individual exon level by developing the ChLoRox-Seq pipeline (Fig. 5, A). Using ChLoRox-Seq, we identified a pool of oxidation-susceptible mRNAs in a cellular model of the human lung epithelium exposed to various environmental oxidative stress conditions. We further investigated these regions of elevated mRNA oxidation to uncover novel characteristics of oxidized RNA regions over non-oxidized RNA regions.

We first demonstrated the specificity of the optimized biotin labeling reaction for targeting 8-oxoG residues compared to unmodified canonical RNA nucleotides (Fig. 1, E). This specificity was benchmarked to the specificity of an anti-8-oxoG antibody used in previous RIP-Seq experiments [9,10]. Ultimately, we found that the biotin labeling method demonstrated comparable specificity to the anti-8-oxoG antibody (Fig. 2, C). Interestingly, while both approaches were successfully able to detect elevated abundance of 8-oxoG in RNA samples exposed to *in vitro* oxidation (Fig. 3, B), only the biotin labeling detection method was able to detect elevated 8-oxoG levels to RNA isolated from *in vivo* oxidized *E. coli* cultures (Fig. 3, D). We hypothesize that this discrepancy may highlight the structural dependence of the anti-8-oxoG antibody for substrate recognition. *In vitro* oxidation at levels employed in this study likely results in elevated levels of denaturation of RNA compared to *in vivo* oxidized RNA samples, thereby increasing accessibility of 8-oxoG sites for the anti-8-oxoG antibody to bind. We expect the biotin labeling method to be less dependent on structure for 8-oxoG detection since it relies on redox chemistry and utilizes a small labeling molecule. A small molecule labeling and detection approach additionally provides a notable cost advantage over antibody-based detection approaches. Specific antibodies for RIP-Seq techniques typically cost approximately $200/pulldown replicate. Conversely, the amount of EZ-link^TM^ Amine-PEG2-Biotin required to perform a similar pulldown technique costs less than $1/pulldown replicate. This reduced cost dramatically eases the financial burden of sequencing experiments and subsequently will enable larger-sized pulldown experiments to be pursued.

### ChLoRox-Seq pipeline allows for functional analysis of oxidation-enriched mRNAs

Having adapted a method developed for DNA [32] to specifically functionalize 8-oxoG sites in RNA with biotin, we further applied this approach to detect the distribution of 8-oxoG accumulation in a transcriptome-wide manner using an affinity capture, pulldown, and sequencing approach, ChLoRox-Seq (Fig. 5, A). We anticipated that this application would help to elucidate factors that bias susceptibility to oxidation of mRNA transcripts across different environmental exposure conditions to human cells. We utilized three different bioinformatics methods to evaluate the pulldown sequencing data and determine regions of elevated RNA oxidation: DESeq2 [48] with individual exon binning, exomePeak2 [49] with GC bias correction, and exomePeak2 without GC bias correction. Ultimately, we observed drastic differences in the regions we identified as oxidized (log2FoldChange > 1, FDR < 0.05) using each method (Table 2). The drastic differences observed were believed to be partly related to the assumptions made in the exomePeak2 pipeline. Since exomePeak2 was originally designed to identify m^6^A peaks in RIP-Seq data [49], it prioritizes identifying distinct regions of enrichment along a given transcript in the enriched pulldown fraction. This approach is effective for analyzing m^6^A distributions, as m^6^A is known to have sequence specificity and occur within a distinct DRACH consensus sequence motif [28,51]. 8-oxoG, having no established consensus motif or writer enzyme in mRNA, is less likely to occur in distinct regions within a transcript. Instead, we believed that 8-oxoG accumulates broadly (i.e. not in distinct regions) within an RNA. This characteristic would not necessarily result in the formation of distinct read “peaks” within a transcript in the enriched fraction of the sequencing data, as is seen in motif-specific modifications such as m^6^A. As a result, we chose to pursue the DESeq2 with exon binning strategy to evaluate regions of elevated mRNA oxidation. This approach evaluates the enrichment of pulldown reads within an exon and is not influenced by the relative enrichment of adjacent exon regions.

Using the DESeq2 with exon binning approach, we identified thousands of regions enriched in the oxidized pool of the RNA-Seq data across different exposure conditions (Fig. 5, B). Interestingly, upon molecular function GO enrichment analysis with ShinyGO version 0.80 [50], we discovered that gene clusters related to actin binding (GO:0003779) and cytoskeleton protein binding (GO:0008092) are upregulated in the oxidized pool under conditions of PM stress (Fig. 5, E). Prior PM exposure work from our lab has demonstrated that similar particulate matter exposure conditions facilitate drastic changes to cellular morphology [44]. Our findings, which suggest elevated oxidation of mRNAs involved in maintaining cellular structure, support the idea that improper expression of oxidized cytoskeleton protein transcripts may contribute significantly to the previously observed morphological alterations upon exposure to PM mixtures. We also observed an enrichment in transcription factor binding (GO:0008314), histone deacetylase activity (GO:0004407), and chromatin binding (GO:0003682) molecular function pathways for genes enriched in the oxidized pool unique to PM exposure conditions, suggesting that these pathways are also implicated in cellular response to PM stress. Similar chromatin-involved pathways were previously found to be enriched in oxidized mRNAs using a RIP-Seq approach to study mRNA oxidation in BEAS-2B cells exposed to gaseous formaldehyde [10], further indicating consistency in RNA oxidation patterns under environmentally induced stress.

### Analysis of the ChLoRox-Seq oxidized RNA pool identifies intrinsic features of oxidized mRNAs

We next investigated the intrinsic genetic traits of the oxidized mRNA pool to identify potential factors that bias mRNA oxidation susceptibility; we refined our investigation to the highest confidence by only considering the mRNA transcripts enriched in the oxidized pool across all exposure conditions (735) (Fig. 5, C). Using general transcript metadata obtained from ShinyGO version 0.80 [50], we discovered that transcripts enriched in the oxidized pool were longer when compared to all the mRNAs detected in the general (Input) pool (Fig. 6, A). This elevated transcript length resulted from longer CDS and 5’ UTR regions (Fig. 6, B-C), as no difference was observed for 3’ UTR length (Fig. 6, D). An increase in the length of 5’ UTR and CDS regions for oxidized pool enriched mRNAs might indicate that these mRNA regions are more susceptible to oxidation. Additionally, we found that oxidized pool enriched mRNAs generally had an elevated number of transcript exons (Fig 6, E).

In looking at the individual nucleotide compositions of oxidized pool enriched exons conserved across all exposure conditions (852) (Supp. Fig. S3, B), we identified that these exons were enriched in G and C nucleotides and de-enriched in A and U nucleotides (Fig. 7, A), relative to the general (Input) pool. This result supports previous work, which identified the enrichment of 8-oxoG in DNA promoter regions harboring GC-rich transcription factor binding sites [33]. Additional analysis into the oxidized pool enriched exons revealed that they were elevated in length compared to exons in the general (Input) pool (Fig. 7, B). When we further reduced our Input population to include only exons within genes that contained an oxidized pool enriched exon, we found similar trends with respect to exon nucleotide composition and length (Fig. 7, C-D). These results suggest that regions of RNA oxidation tend to occur within exons that are uniquely long and elevated in G content relative to other exons within the same transcript.

### Limitations

We acknowledge several limitations of ChLoRox-Seq. Foremost, we observed a high degree of replicate variability amongst pulldown fraction samples (Supp. Fig. S2), particularly in media only CTRL samples (Supp. Fig. S2, B). As a result, we recommend performing a minimum of three biological replicates for each experimental condition evaluated using ChLoRox-Seq. Additionally, while this method allows for the identification of a pool of mRNA sequences that are enriched in 8-oxoG, the phenotypic impact of mRNA oxidation on specific mRNA sequences remains unclear. Future work investigating the mechanistic impact of mRNA oxidation on specific transcripts (i.e. protein misfolding, ribosomal stalling, etc.) is required to connect the oxidation enrichment fold changed quantified using ChLoRox-Seq with phenotypic observations of cellular stress response. Lastly, given the lack of experimentally confirmed mRNAs with heightened propensity for the accumulation of 8-oxoG in the literature, and having not orthogonally confirmed oxidation of individual mRNA transcripts identified using ChLoRox-Seq, we acknowledge that a portion of the oxidized mRNAs identified using ChLoRox-Seq may be false positives resulting from biases inherent to next generation sequencing based analyses, off-target capture, etc.

### Conclusion

The exon-level resolution obtained in this work enabled a deeper investigation into mRNA characteristics that contribute to oxidation accumulation. In this way, ChLoRox-Seq has been presented as an advantageous, readily applicable tool for furthering understanding of dynamic patterns of 8-oxoG accumulation within mRNAs, as well as the mechanisms that might modulate these observed patterns. Taken together, the unique observed features of oxidation-enriched mRNAs presented in this work begin to lay a foundational understanding of the many factors that appear to bias mRNA oxidation susceptibility. Importantly, these observations suggest that most mRNA oxidation events may only occur probabilistically, and that mRNAs that possess certain intrinsic properties (e.g. longer sequence, elevated in G composition, etc.) are more likely to incur oxidation events at elevated rates. These observed sequence traits are unique to the 8-oxoG modification landscape—a stark contrast to other common mRNA modifications (e.g. m^6^A) which are known to be catalyzed by writer enzymes with sequence/structure specificity. However, the clustering of ChLoRox-Seq identified oxidized mRNAs within specific molecular function pathways (Fig. 5, D-E) might point to a larger evolutionary framework that has selected for/against certain mRNAs to act as oxidation sinks. Given the generality of the ChLoRox-Seq approach, future studies can apply this tool to answer additional fundamental questions related to how 8-oxoG accumulation impacts cells under any imposed stress or disease state.

## Materials and Methods

### Cell Culture

BEAS-2B cells (ATCC CRL-9609) were cultured from cryopreserved stocks in T-75 culture flasks according to ATCC recommendations. Cells were seeded at a density of approximately 3,000 cells/cm^2^ and cultured in 23 mL of Airway Epithelial Cell Growth Medium (PromoCell, C-21160) in a humidified incubator (37°C, 5% CO2). A media exchange was performed every 48 hours until the cultures reached 70-80% confluency (approximately 4 days). Cells were then sub-cultured onto 100 mM culture plates, 6-well culture plates, or 96-well culture plates pre-coated with Type 1 collagen (Advanced BioMatrix, Cat #5005) and incubated for 24-48 hours before exposure treatment to enable cell adhesion to the coating matrix.

*Escherichia coli* (*E. coli*) wild-type str. K12 substr. MG1655 was cultured aerobically in Luria-Bertani (LB) broth at 37°C, shaking and on LB-agar plates when required.

### Synthetic RNA Oligonucleotides

8-oxoG modified (N_18_-[8-oxoG]-N_6_) and unmodified (N_24_) RNA oligonucleotides used in this study were purchased as gel purified, lyophilized aliquots from GeneLink. The 8-oxoA modified (N_12_-[8-oxoA]-N_12_) RNA oligonucleotide used in this study was graciously donated from the Resendiz Group at CU Denver.

### RNA Extraction and Purification

RNA purification from BEAS-2B and *E. coli* cells was performed using the DirectZol RNA Miniprep Plus Kit (Zymo Research, R2072) following the manufacturer protocol. Briefly, TRIzol™ Reagent (Invitrogen, 15596026) was added directly to cells immediately after the conclusion of exposure. A DNase I treatment was included during the column clean-up to remove contaminating gDNA from RNA samples. The final elution step of this protocol was performed using RNase-free, N_2_-purged H_2_O in lieu of the provided elution buffer to avoid spurious oxidation of RNA in solution, as described in [9]. RNA purity was confirmed using a NanoDrop 2000 Spectrophotometer (Thermo Scientific) and the concentration of RNA was measured with a Qubit^TM^ 4 Fluorometer (ThermoFisher) using the RNA Broad Range Assay (ThermoFisher, Q10210).

### Biotin Functionalization of 8-oxoG in RNA

Purified RNA was added to an aqueous solution containing 100 mM NaP_i_ Buffer (pH 8.0), 2.5 mM EZ-link^TM^ Amine-PEG2-Biotin (ThermoFisher, 21346), 0.4 U/µL RNase Inhibitor (Murine) (NEB, M0314S). The reaction was started by adding K_2_IrBr_6_ (ThermoFisher, Cat. 11319417) to a concentration of 0.5 mM and allowed to progress for 30 minutes at room temperature (20°C) in the dark. Afterwards, reactions were quenched by the addition of EDTA (Invitrogen, AM9260G) to a concentration of 0.8 mM. RNA was re-purified using the Oligo Clean and Concentrator kit (Zymo Research, D4061) following the manufacturers protocol. Re-purified RNA was quantified with a Qubit 4 Fluorometer (ThermoFisher) using the RNA High Sensitivity Assay (ThermoFisher, Q32855).

### Biotin Dot Blot Assay

Purified, biotin labeled RNA samples were normalized to the same concentration and equal volumes were added to a Amersham^TM^ Hybond^TM^-N+ Western Blotting Membrane (Cytiva, 95038-302). After the samples were allowed to air dry, RNA samples were crosslinked to the membrane using a HL-2000 HybriLinker^TM^ Incubator (UVP, 95003101) set to 120,000 µJ/cm^2^ for 30 seconds. The membrane was blocked in blocking solution (5% (w/v)) dehydrated milk in 1X Tris Buffered Saline + 0.05 % Tween® 20 Detergent (TBS-T)) rocking at 4C in the dark, overnight. The membrane was then briefly rinsed and washed twice in TTBS buffer for 10 minutes at room temperature, rocking. This rinsing and washing process was then repeated using TBS buffer. Afterwards, a 1:2500 dilution of Streptavidin-Horseradish Peroxidase (HRP) Conjugate (ThermoFisher, SA10001) in 1% (w/v) BSA + TBS was added to the membrane and allowed to incubate shaking at 4C in the dark, overnight. The Strep-HRP solution was then removed, and the membrane was similarly subjected to TTBS and TBS washes. The washed membrane was detected using the Clarity Western ECL kit following manufacturer protocol and imaged using a ChemiDoc XRS+ apparatus (BioRad). Pixel density for each dot was quantified using the volume tool in ImageLab^TM^ software (BioRad).

### 8-oxoG Antibody Dot Blot Assay

8-oxoG Antibody Dot Blot Assays were performed on purified, unlabeled RNA samples in the same fashion as the Biotin Dot Blot Assays, with the following exceptions. A 1:2500 dilution of 8-Hydroxyguanine (8-oxo-dG) MAb (Clone 15A3) primary antibody (Bio-techne, 4354-MC-050) in 1% (w/v) BSA + TBS was used instead of the Streptavidin-Horseradish Peroxidase (HRP) Conjugate solution. Additionally, an extra 1-hour room temperature incubation in 1:2500 anti-Mouse IgG-HRP secondary antibody (Promega, W4021) + 1% BSA + TBS was performed prior to Clarity Western ECL detection.

### In vitro oxidation of E. coli total RNA

*In vitro* oxidation of RNA was performed following the protocol outlined in [31]. Briefly, 15 µg of purified total *E. coli* RNA was added to TMN buffer solution (50 mM Tris/HCl pH 7.6, 100 mM NH_4_Cl, 10 mM MgCl_2_, 6 mM *β*-ME) supplemented with 10 µM Fe(*II*)Cl_2_ and H_2_O_2_ ranging from 0-20 mM. This reaction was allowed to proceed for 1 hour at 37°C. Afterwards, RNA was repurified using the Oligo Clean and Concentrator kit (Zymo Research, D4061) and stored at -80C until downstream use.

### In vivo oxidation of E. coli cultures

A range of H2O2 concentrations (0-20 mM) diluted in PBS were added to exponential phase *E. coli* cultures (OD = 0.5-0.6) and incubated at 37C, shaking for 20 minutes. Cells were subsequently harvested via centrifugation at 4,000 rpm, 4°C and immediately flash frozen in liquid nitrogen. Cell pellets were stored at -80°C until use for RNA extractions.

### In vivo oxidation of BEAS-2B cultures

BEAS-2B cells were seeded into individual wells of a 6-well plate at a density of ∼225,000 cells/well and incubated for 24 hours at 37°C, 5% CO2. The growth media was then replaced with fresh BEGM supplemented with 100 µM H2O2 and cells were incubated at 37°C, 5% CO2 for 20 minutes. Afterwards, the growth media was removed and cells were immediately lysed in TRI reagent. RNA was extracted according to the procedure outlined in *RNA Extraction and Purification*.

### Intracellular ROS Detection Assay

BEAS-2B cells were seeded into individual wells of a 96-well plate at a density of ∼10,000 cells/well and incubated for 24 hours at 37°C, 5% CO2. Cells were then loaded with 100 µL of growth media containing 10 µM H_2_DCFDA dye (ThermoFisher) for 30 minutes at 37°C, 5% CO2. The dye was then removed and replaced with 100 µL of fresh growth media (CTRL) or growth media supplemented with 100 µM H2O2 (H2O2), 500 µg/mL urban particulate matter (PM) (NIST, SRM 1648a), or 500 µM formaldehyde (FA). The A493/522 fluorescence was immediately monitored in 5-minute intervals using a BioTek Cytation3 microplate reader over the course of 60 minutes with the cells being maintained at a constant 37°C.

### Preparation of exposure samples for ChLoRox-Seq

BEAS2B cells were seeded onto 100 mM culture plates at a density of 500,000 cells/plate and grown for 48 hours in 22.4 mL of growth media. The media was then exchanged for each plate and replaced with fresh media (CTRL) or media that was supplemented with an exposure agent (100 µM H2O2 or 500 µg/mL urban particulate matter (PM) (NIST, SRM 1648a)). These cultures were incubated for 20 minutes at 37°C, 5% CO2. Afterwards, cells were immediately lysed in TRI reagent and RNA was extracted according to the procedure outlined in *RNA Extraction and Purification*. These samples were used for subsequent ChLoRox-Seq analysis.

### ChLoRox-Seq Pipeline

The quality of total RNA extracts was evaluated on an Agilent BioAnalyzer 2100, with all samples generating a RIN score > 9. Purified RNA was poly(A)-selected using the NEBNext® High Input Poly(A) mRNA Isolation Module (NEB, E3370S). The final elution step of this protocol was performed using RNase-free, N_2_-purged H_2_O in lieu of the provided elution buffer. Afterwards, mRNA samples were fragmented using the NEBNext® Magnesium RNA Fragmentation Module (NEB, E6150S) at 94°C for 4 minutes. Fragmented mRNA was re-purified using the Oligo Clean and Concentrator Kit (Zymo Research, D4061). Following purification, RNA samples were subjected to the biotin labeling reaction and clean-up in accordance with the *Biotin Functionalization of 8-oxoG in RNA* protocol described earlier. 1/10^th^ of the volume of each of the purified RNA samples was isolated to create NGS input libraries (Input). The remaining portion of each sample was used in Streptavidin Dynabead^TM^-mediated pulldowns to isolate biotinylated RNA fragments following manufacturer protocol (ThermoFisher, 65001). Briefly, an equal volume of 2X B&W buffer (10 mM Tris-HCl (pH 7.5), 1 mM EDTA, 2 M NaCl) was added to homogenized Dynabeads™ and mixed by rotation for 5 minutes at room temperature. A magnetic rack was used to separate beads and the supernatant was discarded. This process was repeated twice with 1 mL of 2X B&W buffer for a total of 3 washes. The beads were then washed twice in 1 mL of Solution A (0.1 M NaOH, 0.05 M NaCl) and twice in 1 mL of Solution B (0.1 M NaCl), rotating for 2 minutes each wash. Afterwards, the beads were resuspended in 2X B&W buffer supplemented with 0.4 U/µL RNase Inhibitor (Murine) (NEB, M0314S) to a bead concentration of 5 ug/µL. An equal volume of each RNA sample was then added to the bead mixture and the samples were allowed to rotate at room temperature for 20-30 minutes. Unbound RNA was removed by magnetic separation and the bound fraction was washed 3 times in 1 mL of 1X B&W buffer as previously described. A fourth wash was performed with 200 µL of B&W buffer and the mixture was transferred to a nuclease-free 0.2 mL PCR tube to ensure complete removal of supernatant buffer. To remove bound RNA from the Dynabeads^TM^, samples were suspended in 20 µL of non-ionic, sterile H_2_O and heated in a thermalcycler (BioRad) at a rate of 0.5°C/second from 25°C to 70°C. Samples were incubated at 70°C for 1 second and then slowly cooled back to 25°C at a rate of 0.5°C/second. Immediately afterwards, dissociated RNA fragments were collected from the supernatant solution. These recovered RNA fragments were used to create NGS pulldown libraries (Pulldown).

Library preparation and sequencing was performed by the University of Texas at Austin Genomic Sequencing and Analysis Facility (GSAF). Briefly, libraries were prepared for all samples using the NEB Next Ultra II Directional RNA Library Prep Kit (NEB, E7760S) according to manufacturer protocol. The resulting libraries tagged with unique dual indices were checked for size and quality using the Agilent High Sensitivity DNA Kit (Agilent). Library concentrations were measured using the KAPA SYBR Fast qPCR kit (Roche) and loaded for sequencing on the NovaSeq 6000 instrument (Illumina). Libraries were run in paired-end, 100 bp mode across two lanes to a minimum depth of 10 million reads/sample for pulldown fraction samples and minimum depth of 20 million reads/sample for Input fraction samples. Raw, unfiltered reads were aligned to the UCSC reference genome, hg38.p14, using the STAR aligner version 2.7.0d with default settings [52]. SAM files were sorted into BAM files using the SAMtools sort function and merged across sequencing lanes using the SAMtools merge function [53].

Aligned BAM files and the UCSC hg38.refGene.gtf file were provided as input for the exomePeak2 [49] analysis pipeline to elucidate differential enrichment of reads around modification locations. The UCSC hg38.p14.fa genome file and was supplied as additional input for default exomePeak2 analysis to correct for genomic GC bias. The genome file was not supplied as input when running exomePeak2 without GC bias correction analysis. Oxidation peak regions were determined from the program output by filtering for RPM_input_>1, log2FoldChange (Pulldown/Input) >1, and FDR < 0.05.

For DESeq2 [48] with exon binning analysis, aligned BAM files were binned to UCSC hg38.refGene.gtf exon features using HT-Seq [54] (htseq-count -f bam -s reverse -m union -t exon -r pos -i gene_id -i exon_number -a 30 --additional-attr=gene_name --additional-attr=exon_number --nonunique=random). The resultant unnormalized count files were filtered for a mean average read count per feature of greater than 10 across input fraction samples. These filtered count files were subsequently used as input for DESeq2 [48] with the fitType set to ‘local’. Oxidized exon regions were classified as having a log2FoldChange (Pulldown/Input) > 1 and p_adj_ < 0.05. Exon specific sequence data was obtained using the Hsapiens.UCSC.hg38 genome from the BSgenome package [55] in R in combination with the UCSC hg38.refGene.gtf file.

Data was visualized using the Tidyverse [56] and cowplot [57] packages in R. All output tables from DESeq2 and exomePeak2 bioinformatic analysis pipelines are provided in supplemental tables S1-S18.

## Data Availability Statement

The data that support this study are available from the corresponding author upon reasonable request. All ChLoRox-Seq FASTQ files generated during this study are available for download on the NCBI Sequence Read Archive (SRA) database under BioProject accession number PRJNA1120946.

## Disclosure Statement

No potential conflicts of interest are reported by the authors.

## Funding Statement

This work was supported by grants from the National Science Foundation [grant numbers MCB-2218477 to M.R.B, P. J. S., and L.M.C., DGE-2137420 to M.R.B.).

## Acknowledgements

The authors acknowledge the Texas Advanced Computing Center (TACC) at The University of Texas at Austin for providing high performance computing resources that have contributed to the research results reported within this paper. URL: http://www.tacc.utexas.edu. We also acknowledge the University of Texas Genomic Sequencing and Analysis Facility (GSAF) for library preparation and RNA sequencing. The authors also acknowledge the use of <u>Biorender.com</u> for graphical figure creation.

## Notes

### Competing Interest Statement

The authors have declared no competing interest.

