## Supplemental_Figures for "Optimized chemical labeling method for isolation of 8-oxoG-modified RNA, ChLoRox-Seq, identifies mRNAs enriched in oxidation and transcriptome-wide distribution biases of oxidation events post environmental stress"

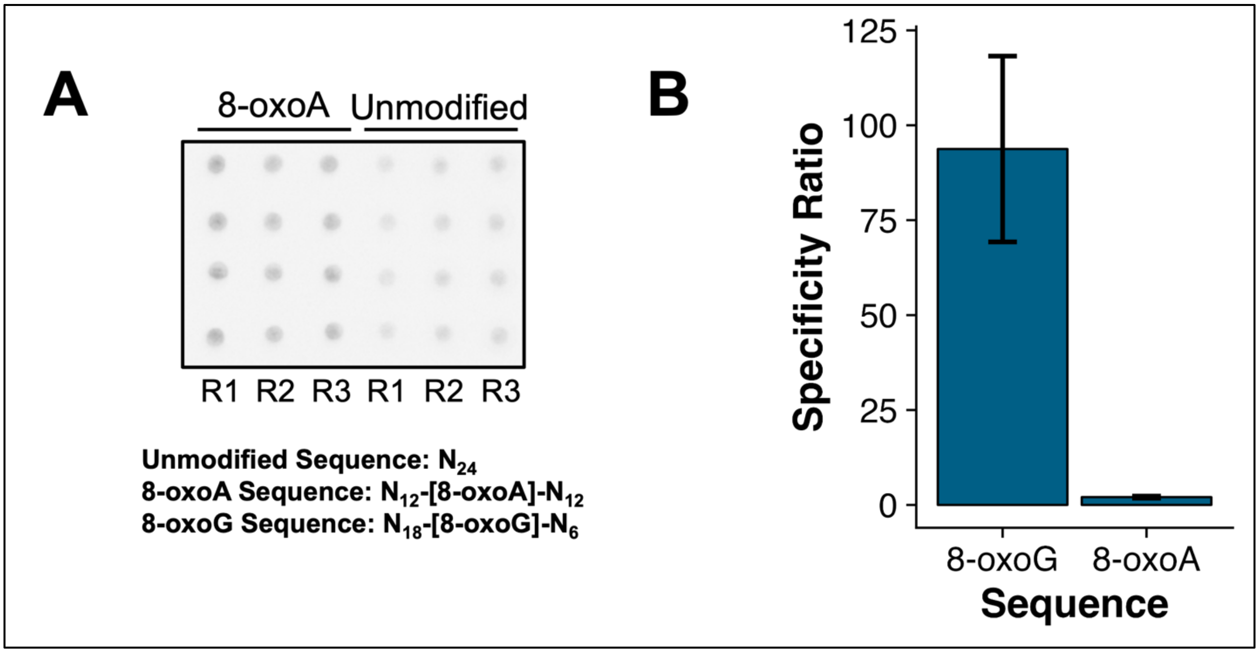


***Supplemental Figure S1.*** The biotin labeling reaction preferentially labels 8-oxoG modifications over 8-oxoA modifications. (A) (Top) Dot blot image of biotin labeling reaction products for 8-oxoA modified RNA versus unmodified RNA. (Bottom) Sequences of synthetic RNA oligonucleotides used in dot blot specificity analysis. (B) Bar graph of quantified specificity ratios for biotin labeling reaction on 8-oxoG-modified versus unmodified RNA templates and 8-oxoA-modified versus unmodified RNA templates.

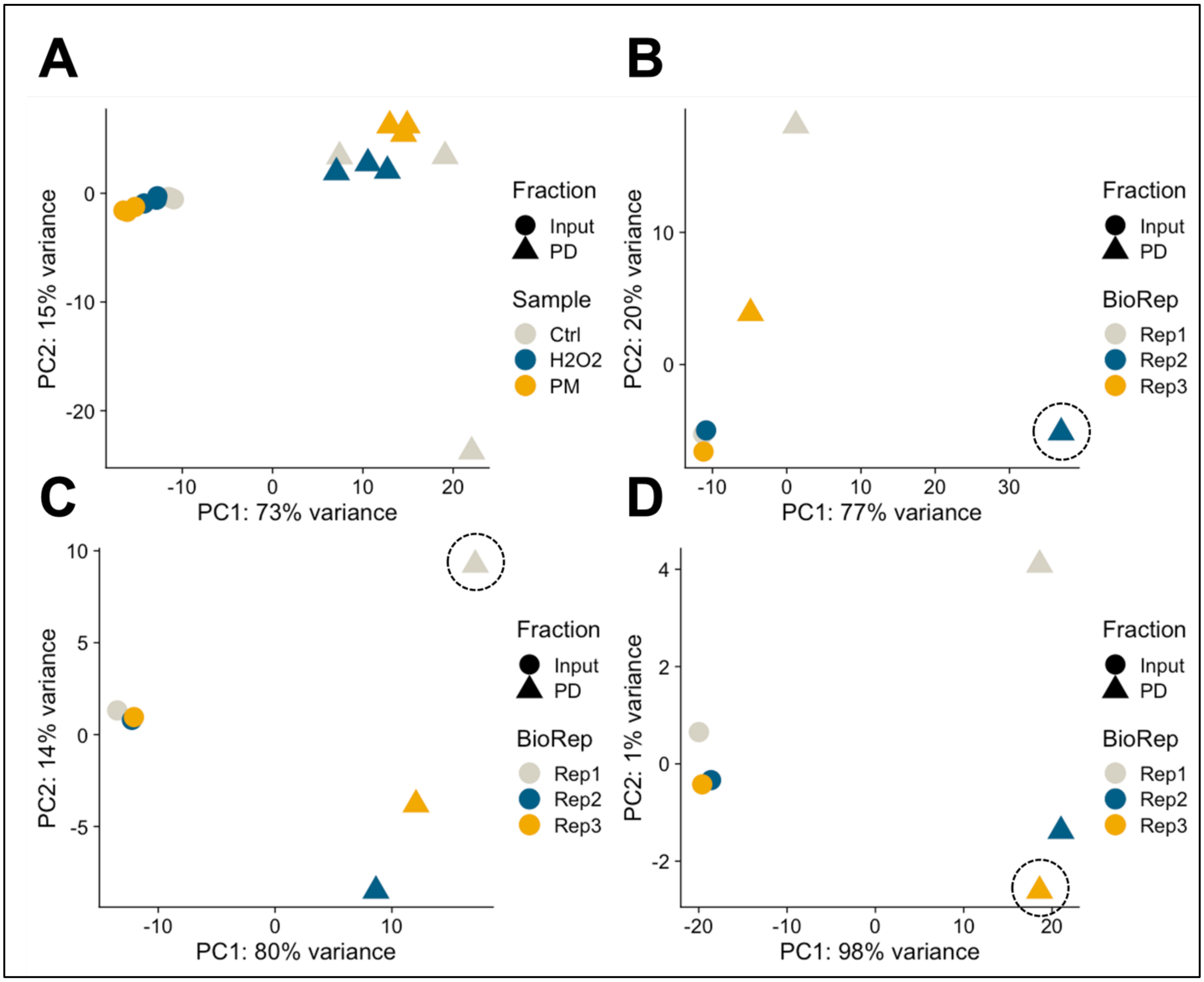


***Supplemental Figure S2.*** Principal component analysis (PCA) depicts general clustering of RNA-seq samples from input (Input) and pulldown (PD) fractions for each tested exposure condition. (A) PCA plot of samples from all experimental conditions annotated by Fraction and Sample. (B-D) PCA plot of media only (CTRL) (B), 100 µM H2O2 (H2O2) (C), and 500 µg/mL PM (PM) (D) samples annotated by Fraction and BioRep. Black dashed circles indicate the pulldown replicate that was dropped during comparative analysis between DE-Seq2 and exomePeak2 bioinformatic approaches (Table 2).

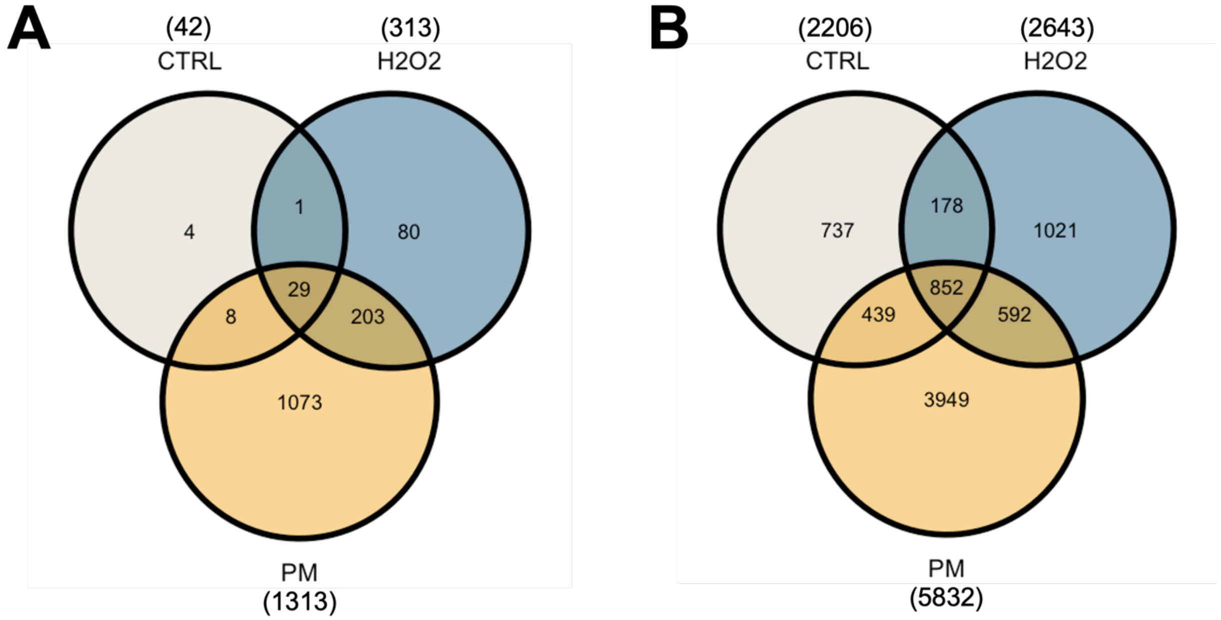


***Supplemental Figure S3.*** Comparison of overlapping regions of RNA oxidation across exposure conditions. (A) Venn diagram showing number of overlaps of oxidized region containing genes across exposure conditions identified by all three bioinformatics processing approaches (DE-Seq2 with exon binning, exomePeak2 with GC bias correction, and exomePeak2 without GC bias correction). (B) Venn diagram showing number of overlaps of oxidized exon regions conserved across exposure conditions when evaluating using the DE-Seq2 with individual exon binning bioinformatic processing approach.

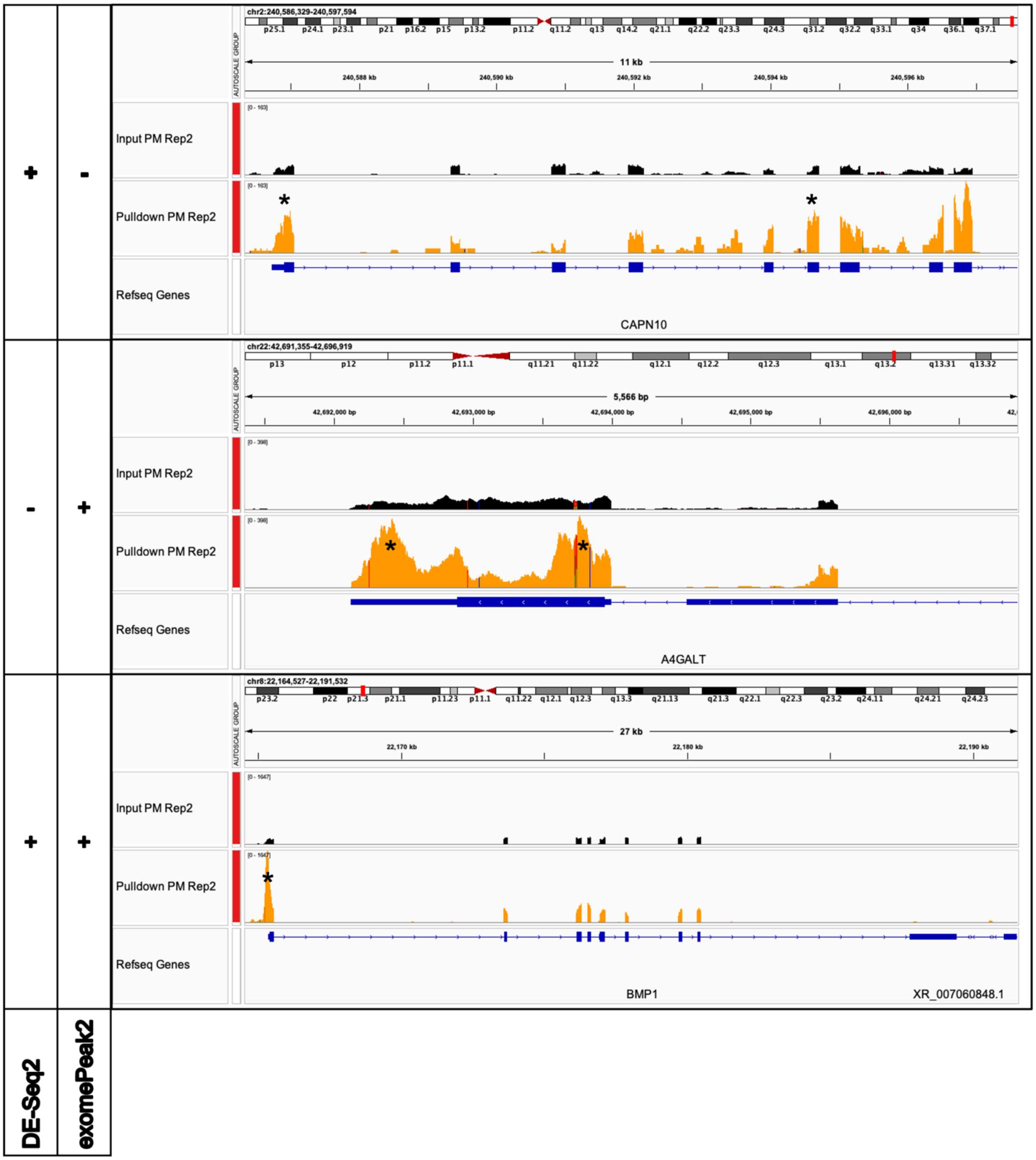


**Supplemental Figure S4.** IGV browser screenshot of example regions of predicted RNA oxidation as detected by the DE-Seq2 with individual exon binning approach and the exomePeak2 with GC bias correction approach for PM exposure replicate 2. (+) Indicates a positive hit for RNA oxidation, (-) indicates a negative hit for RNA oxidation, (*) indicates the bioinformatically predicted location of RNA oxidation.

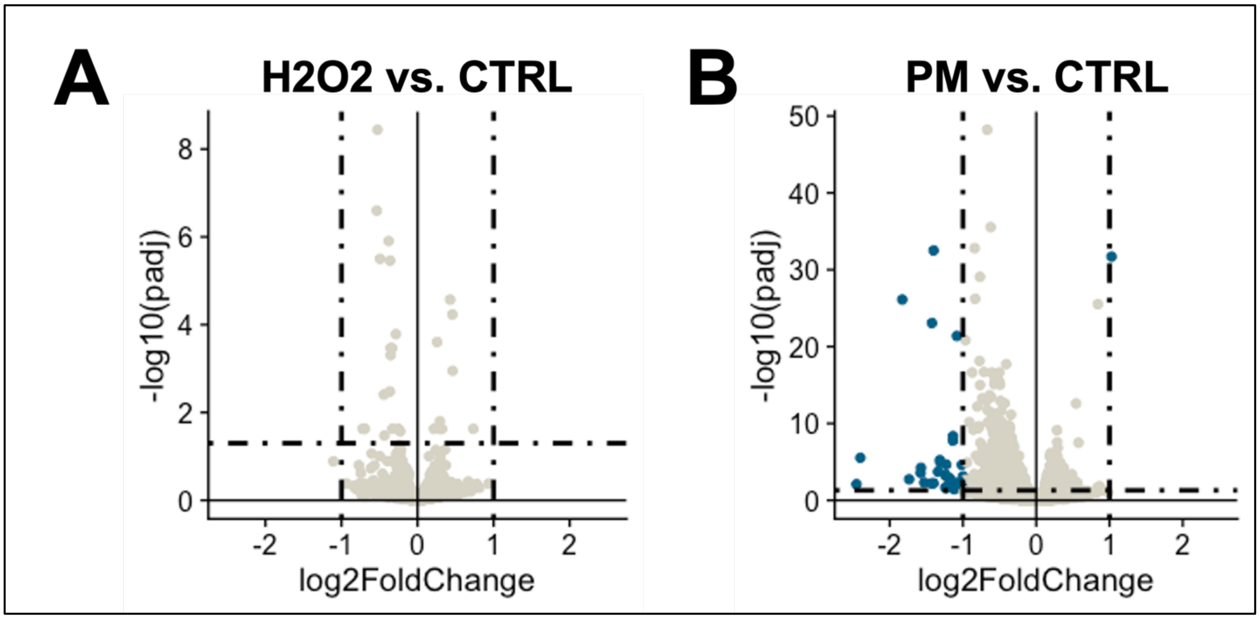


**Supplemental Figure S5.** Exposure conditions used in biotin pulldown and sequencing experiments do not trigger a large-scale transcriptional response. (A) Volcano plot depicting differential gene expression in 100 µM H2O2 exposed samples versus media only CTRL samples. (B) Volcano plot depicting differential gene expression in 500 µg/mL H2O2 exposed samples versus media only CTRL samples. Blue dots indicate genes that pass the established thresholds for differential expression: | log2FoldChange | > 1 AND $p_{adj} < 0.05$.
